# Multihospital expansion of vancomycin-resistant *Enterococcus faecium* ST117-CT7799 and transmission of linear plasmids co-carrying *vanA* and linezolid resistance genes, Comunitat Valenciana, Spain (2022–2024)

**DOI:** 10.64898/2026.06.01.729239

**Authors:** Carlos Valiente-Mullor, Lidia Ruiz-Roldán, Ana C. Almeida-Santos, Regina Antoni, Alejandro Sanz-Carbonell, Sara Arnal, Susana Sabater, Ignacio Torres, Eva Gonzalez-Barbera, Inmaculada Vidal, Nuria Tormo, Rafael Medina-González, Carla Novais, Luisa Peixe, Ana R. Freitas, Fernando González-Candelas

**Affiliations:** Joint Research Unit Infection and Public Health FISABIO-Univ. Valencia, Institute for Integrative Systems Biology (I2SysBio, CSIC-UV). Valencia, Spain; UCIBIO, Unidade de Ciências Biomoleculares Aplicadas, Faculdade de Farmácia, Universidade do Porto, Porto, Portugal; Laboratório Associado i4HB, Instituto para a Saúde e a Bioeconomia, Faculdade de Farmácia, Universidade do Porto, Porto, Portugal; Servicio de Microbiología, Hospital Clínico Universitario Lozano Blesa, Zaragoza, Spain; Servicio de Microbiología, Consorcio Hospital General Universitario de Castelló, Castelló, Spain; Servicio de Microbiología, Hospital Clínico Universitario, Valencia, Spain; Servicio de Microbiología, Hospital Universitario y Politécnico La Fe, Valencia, Spain; Servicio de Microbiología, Hospital Francesc de Borja, Gandía, Spain; Servicio de Microbiología, Consorcio Hospital General Universitario de Valencia, Valencia, Spain; UCIBIO, Unidade de Ciências Biomoleculares Aplicadas, Instituto Universitário de Ciências da Saúde (1H-TOXRUN, IUCS-CESPU), Gandra, Portugal; CIBER in Epidemiology and Public Health, ISCIII, Madrid, Spain

**Keywords:** Enterococci, genomics, vancomycin resistance, linezolid resistance, linear plasmids, bacteriocins, bac43

## Abstract

**Background:** Vancomycin-resistant *Enterococcus faecium* (VREfm) is a WHO priority pathogen. In the Comunitat Valenciana (CV), Spain, VREfm prevalence has increased since 2022. We characterized the population structure, transmission patterns and resistance determinants of VREfm across eight hospitals (2021-2024).

**Methods:** Eight hospitals reported 870 VREfm cases during 2021–2024. We sequenced 254 VREfm isolates using WGS (Illumina) and inferred relatedness by MLST, cgMLST, and core-genome SNP analyses. Acquired antimicrobial resistance (AMR) genes, bacteriocins, plasmid replicases and putative virulence markers (PVMs) were identified *in silico*. Thirty-eight representatives underwent Nanopore long-read sequencing and hybrid assembly to resolve plasmids.

**Results:** The predominant vancomycin-resistance genotype was *vanB* (62%; six hospitals), followed by *vanA* (19%; five hospitals) and *vanA*+*vanB* (17%; five hospitals). Eight sequence types (ST) and 15 clonal complexes (CT) were identified, grouped into seven main phylogenetic clades with close relatedness to publicly available genomes from other regions of Spain and Europe. A single lineage, ST117-CT7799, accounted for 198/254 (78%) isolates, persisted during 2022-2024 across seven hospitals and was enriched in the bacteriocin gene bac43 (or T8) (84%). Linezolid-resistance genes *optrA* and *cfr*(D) were present in 34% of isolates, most of which also carried *vanA* (32%). Hybrid assemblies revealed a diverse plasmidome including RepA_N megaplasmids with PVMs and AMR genes, and small Rep3-like plasmids harbouring bacteriocins (*bac43*, *bac51*, *bacAS9*); a 6 kb repA_pB82 plasmid was *bac43*-positive in 37/38 (97%) fully resolved carriers. Eight strains across four hospitals carried identical linear plasmids co-harbouring *vanA*, *optrA*, *cfrD* genes within a widespread repUS78_pZY2 background, consistent with inter-lineage plasmid transmission.

**Conclusions:** This first comprehensive genomic analysis of VREfm in CV indicates extensive inter-hospital spread dominated by expansion of ST117-CT7799 and highlights plasmid-mediated convergence of vancomycin and linezolid resistance via linear plasmids. Strengthened infection prevention and genomic surveillance, including long-read sequencing to track linear plasmids, are warranted.

## INTRODUCTION

*Enterococcus faecium*, a natural gut commensal bacterium, has emerged as a major opportunistic pathogen [1]. Nosocomial infections caused by *E. faecium* have become a global concern, due to its persistence and adaptability to the healthcare environment, intrinsic resistance to multiple antibiotics (e.g., aminoglycosides and cephalosporins), and the acquisition and spread of novel virulence and antimicrobial resistance genes (ARG) [2]. Among hospital-associated *E. faecium* [3], ST80 and ST117 are two of the emergent lineages of concern, showing persistence in the hospital setting and responsible for nosocomial outbreaks in several European countries including Spain [4–7].

More concerning is vancomycin-resistant *E. faecium* (VREfm), which is included in the WHO priority list of antibiotic-resistant bacteria as a high-risk pathogen, due to limited treatment options and its high propensity to acquire antimicrobial resistance [8]. In Europe, the prevalence of VREfm associated bloodstream infections has been on the rise since 2000, although marked differences in geographical distribution are observed [6,9,10]. Despite the prevalence of VREfm being lower in Spain than in other EU/EEA countries (2.6% of invasive isolates in 2024 vs. an average of 16.5%), a significant increase has been observed since 2020, peaking in 2023 at 4% [10]. This follows a decline between 2015-2019, from 2.5% to 1.2% [11]. Linezolid remains one of the few therapeutic options available for treating VREfm infections [12]. Although the overall incidence of linezolid-resistant *E. faecium* (LREfm) and linezolid-resistant VREfm (LVREfm) remains low, and systematic surveillance is lacking in most countries [7,13], the growing number of reported infections and outbreaks involving LREfm is concerning [14,15].

In recent years, the incidence of VREfm has increased in the Comunitat Valenciana (CV) region of Spain, with cases reported independently across different healthcare settings. However, no comprehensive analysis has yet been conducted to characterize the clonal relationships and mobilome of the circulating lineages. Therefore, our aim was to conduct a retrospective genomic analysis (2021-2024) of more than 200 clinical *E. faecium* isolates from eight hospitals in the CV region, with a focus on the epidemiology of vancomycin and linezolid resistance, as well as the characterization of other adaptive traits, including plasmids, virulence markers, and bacteriocins.

## MATERIALS AND METHODS

### Bacterial collection and study design

Between 2021 and 2024, a total of 264 VREfm were recovered from eight hospitals in the CV region, Spain. The study included four tertiary care hospitals (521-1000 beds) located in the provincial capitals - Castellón (HGUC) and Valencia (HLAFE, HCUV, HGRAL)- and their associated secondary hospitals (HMAG, HSAG, HFBG, HLAX), ranging from 128 to 285 beds. An initial increase in VREfm cases was detected in 2022 at HGUC and HMAG (Castellón, Spain), prompting the selection of 32 isolates (2022-2023) for short- and long-read sequencing. Concurrently, a similar rise in VREfm incidence was observed across additional healthcare centres in the region, motivating to expand the study to a broader regional scale. Of the 1110 VREfm cases reported in the CV region between January 2022 and September 2024, 212 additional isolates, approximately 20%, were longitudinally selected for short-read sequencing from the five hospitals with the highest VREfm prevalence (HGUC, HLAFE, HCUV, HFBG, HSAG). Of these, nine representatives from distinct lineages were selected for additional long-read sequencing. Furthermore, to ensure broader regional and temporal representation, short-reads of 12 isolates (2022-2024) from remaining lower-prevalence centres (HGRAL, HLAX) were included, together with eight isolates from 2021 from a previous outbreak at HFBG (Table S1, Table S2).

### Whole genome sequencing and data analysis

Strains were cultivated on blood agar plates. A single colony was inoculated on broth BHI tubes and incubated 37°C overnight with agitation. Genomic DNA of bacterial isolates were extracted from 1 mL of culture using either Maxwell RSC Blood DNA kit (Promega) on Maxwell RSC extractor or easyMAG (Biomérieux) extractor (Table S2). DNA was quantified using Qubit 1X dsDNA BR (Invitrogen), quality and contamination were assessed with NanoDrop (Thermo Fisher Scientific) and fragment sizes were determined with TapeStation for genomic DNA (Agilent Technologies).

For short-read sequencing, libraries were prepared using the Illumina DNA Prep kit (Illumina) and the isolates were sequenced on a NextSeq2000 system (Illumina) using the NextSeq P2 kit (600-cycle, 2×300). Short reads were analysed with EPIBAC v1.0.0 pipeline (https://github.com/EpiMol/epibac/) and 14 isolates out of 264 were discarded due to contamination or sequencing failure, generating a final number of 250 short-read sequences. Isolates were further screened for ARGs [ABRicate (https://github.com/tseemann/abricate)] and 23S rRNA mutations associated with linezolid resistance (LRE-finder [16]). We determined the sequence type (ST) with MLST 2.23.0 (https://github.com/tseemann/mlst), using the two typing schemes available for *E. faecium*, hereinafter referred to as the ‘Homan scheme’ [17] or ‘Bezdíček scheme’ [18], respectively. Core genome MLST (cgMLST) and mutations associated with daptomycin resistance were determined using Ridom-SeqSphere+ v11.1.1 (Münster, Germany). Assemblies were screened for bacteriocins and putative virulence markers (PVMs) with BLAST v2.12.0+ [19] (non-redundant hits with >80% identity and >80% coverage) using, respectively, a custom database of *Bacillota* bacteriocin genes [20] and VirulenceFinder 2.0 database [21]. Replicase genes were searched in all assembled genomes using BLAST with PlasmidFinder v2.1 database [22], including novel *rep* gene sequences recently reported [23].

For long-read sequencing, libraries were prepared using the Native Barcoding Kit (SQK-NBD114.24, Oxford Nanopore Technologies [ONT]) and a total of 41 isolates were sequenced on a MinION Mk1C system (ONT) using a R10.4.1 flowcell. After basecalling [DORADO v0.7.3 (https://github.com/nanoporetech/dorado)] and QC [Kraken2 v2.1.3 [24], Filtlong v0.2.1 (https://github.com/rrwick/Filtlong), Porechop v0.2.4 (https://github.com/rrwick/Porechop)], hybrid assemblies were obtained with Hybracter [25] for 34 isolates and four isolates were assembled using only long-reads with Autocycler v0.2.0 [26], as the corresponding short-read sequencing could not be carried out. Three isolates out of 41 were discarded due to contamination, generating a final number of 38 hybrid/ONT assemblies. Additionally, ARGs, bacteriocins, PVMs and plasmid replicases were characterized as previously outlined.

### Identification of related genomes

*E. faecium* whole genome assemblies (n=40,572) were downloaded from AllTheBacteria (ATB) 0.2 incremental release August 2024 [27]. STs were determined as in the previous section. Short reads were compared to the respective subset of ATB assemblies with the same ST (Homan scheme) using Mash v2.3 [28]. The short reads of the closest genome to each isolate and all ST117-ST143 (Bezdíček scheme) genomes in the ATB were downloaded, respectively, from the NCBI-SRA database (Table S3). Additionally, we included short reads of two close isolates (ST117-ST143, ST80-ST152) from a study in a geographically close region [5] that were absent in ATB. Short-read analysis was performed as outlined in the previous section.

### Phylogenomic analysis

Only short reads were mapped with Snippy v4.6.0 (https://github.com/tseemann/snippy) using the hybrid assembly of a ST117-CT7799 isolate from this collection (EPI00100, 2022) as the reference. ≥75% of reads supporting the alternative allele were required for variant calling. Mapping statistics were calculated using bash and samtools v1.20 [29] from resulting BAM files and consensus sequences. Three sequences with coverage <70%, average depth <30 and >10^6^ N bases or gaps were removed from subsequent analysis. Three core genome consensus sequence alignments (including variant and invariant sites) were generated with *snippy-core*, including (a) all the mapped sequences and their closest ATB genomes, (b) only ST80 and ST117 mapped sequences, and their closest ATB genomes, and (c) only ST117-ST143 sequences in this study and ATB.

Sites in <95% of sequences were removed from the alignments with Core-SNP-Filter v0.2.0 [30]. SNP distances between sequences were calculated with snp-dists v0.8.2 (https://github.com/tseemann/snp-dists). Maximum likelihood (ML) phylogenies were built using IQ-Tree2 v2.3.4.cmaple [31], the GTR+G+I substitution model and 1000 ultrafast bootstrap replicates.

### Characterization of *vanA*-carrying linear plasmids

Complete plasmid sequences carrying the *vanA* operon were grouped using pling v2.0.0 [32] according to their containment distance (i.e., proportion of the smaller plasmid not contained in the larger one) and rearrangement (DCJ-indel) distance (i.e., minimal number of rearrangements needed to convert one plasmid into the other). Containment distance threshold was set to 0.3 in order to identify potential transmission events. Comparison of plasmids was visualized with GenoFig v1.1 [33]. To confirm linear topology *in silico*, contig ends of *vanA* plasmids were investigated as in Almeida-Santos et al. (2025) [34] and McCraken et al. (2026) [35].

## RESULTS

### Description of VREfm isolates

A total of 264 VREfm and LREfm strains isolated between 2021 and 2024 in the eight hospitals included in the study were analysed. After initial quality control of the sequencing reads, ten isolates were removed from the collection generating a final number of 254 isolates for this study (Table 1).

**Table 1.**
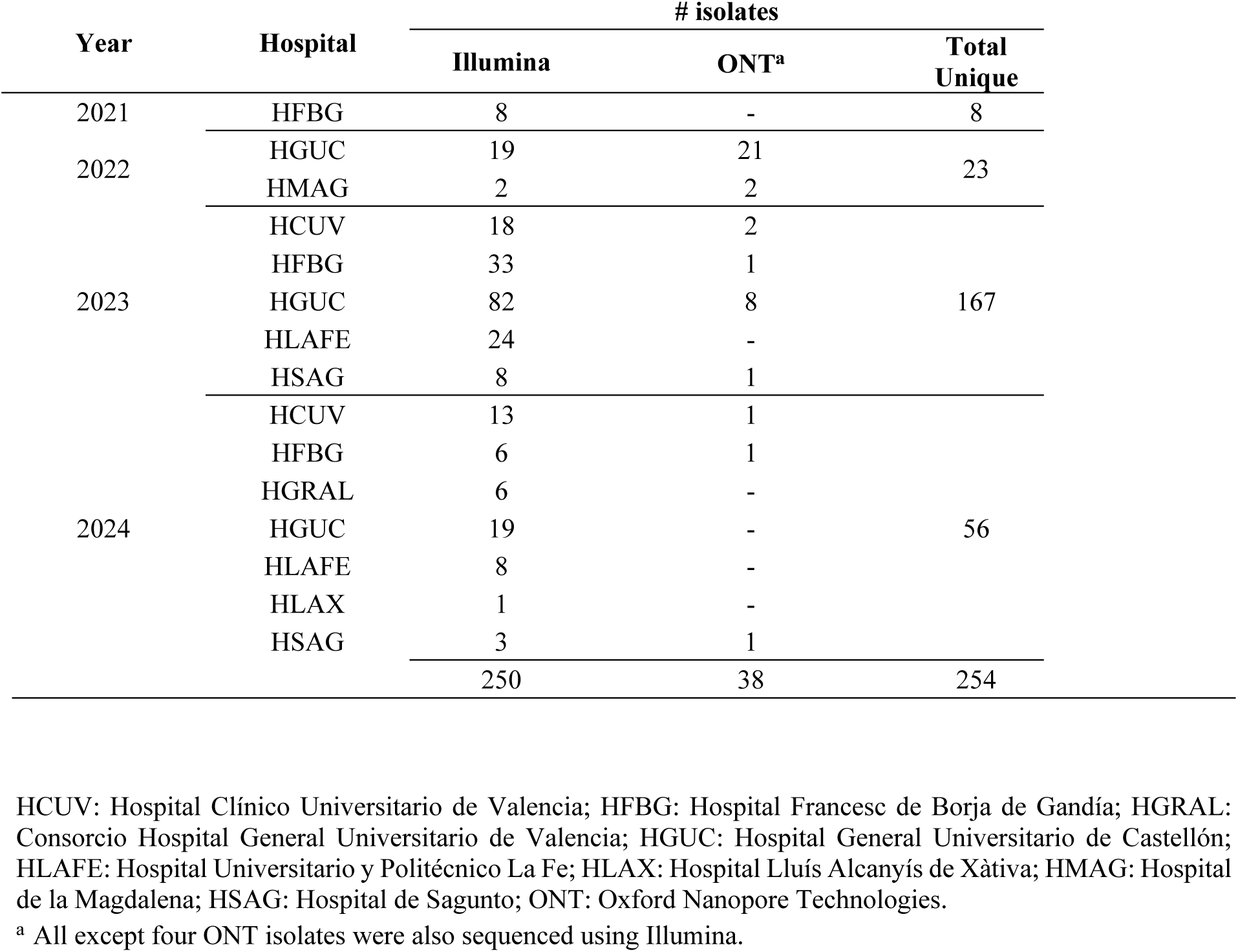
*E. faecium* isolates included in the study by year, hospital and sequencing platform.

Eight STs (Homan scheme) were identified, with ST117 by far the most prevalent (n=221, 87.0%), followed by ST80 (n=20, 7.8%), which was detected exclusively in five of the six hospitals in Valencia province (Figure 1; Table 2; Table S2). Fifteen complex types (CTs) were characterized: six included ST117 isolates, and four corresponded to ST80. The dominant clone ST117-CT7799 (n=197, 77.5%) was detected in all hospitals except HLAX (n=1) during 2022-2024, followed by ST80-CT8599 (n=13, 5.1%) (Figure 1, Table 2, Table S2). ST1886-CT2290 (n=5, 1.9%) strains were isolated from a single hospital in 2021 and were phylogenetically unrelated to the remaining cases. The five singleton ST/CTs were also unrelated (Figure S1).

**Figure 1.**
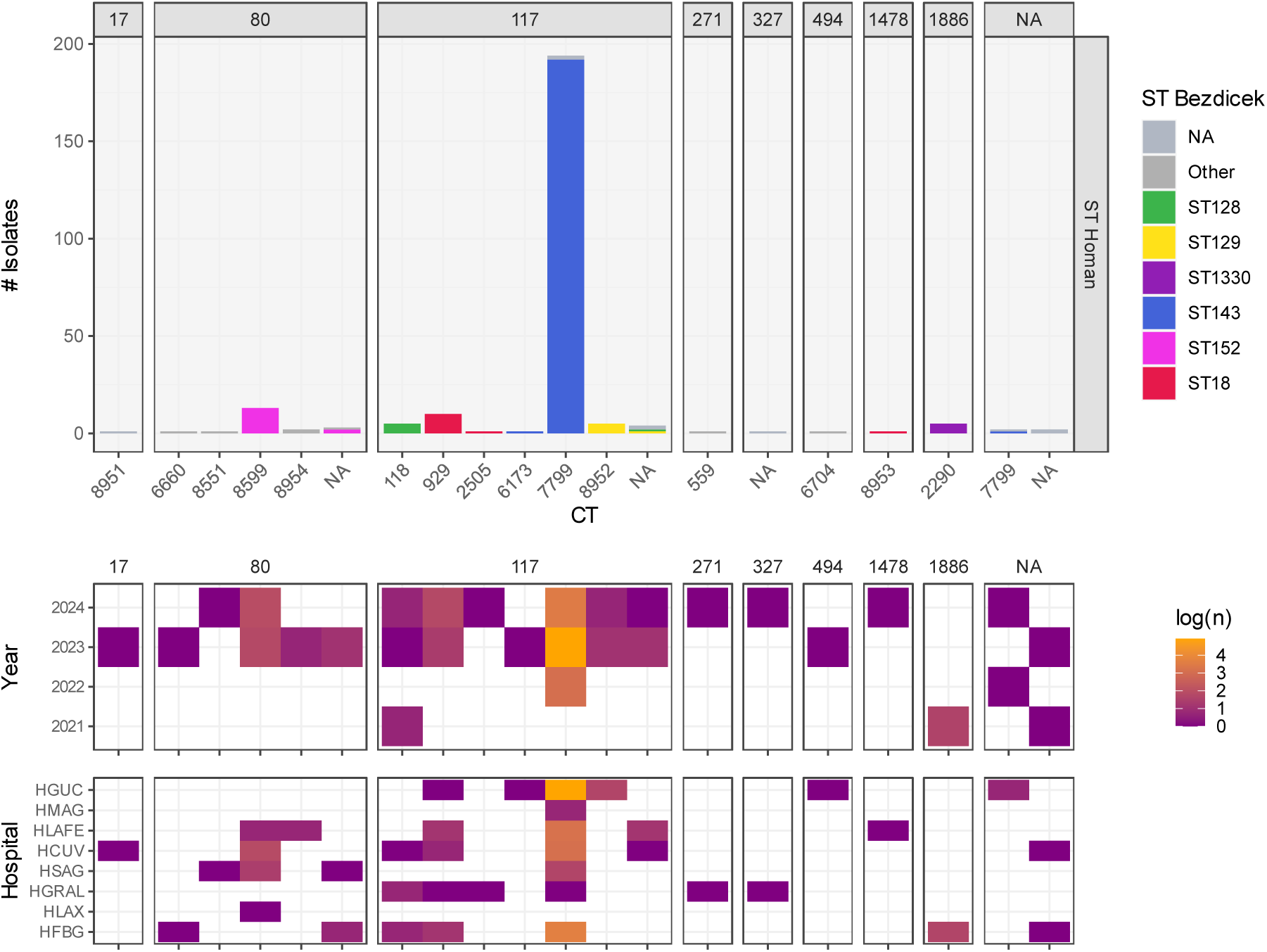
Frequency of sequence and complex types by hospital and year of isolation. STs (Homan scheme) of the 254 isolates analysed are shown in panel labels. Their corresponding CTs (X-axis) and STs (Bezdíček scheme) (stacked bar colour) are shown within each panel.

**Table 2.** Frequency of STs/CTs, and vancomycin and linezolid resistance genotypes.

| ST / cgMLST |  |  |  |  |  | Vancomycin resistance |  |  |  | Linezolid resistance |  |  |  |  |
| --- | --- | --- | --- | --- | --- | --- | --- | --- | --- | --- | --- | --- | --- | --- |
| Homan scheme | Bezdiček scheme | CT | Year | # Isolates | Total | *vanA* | *vanB* | *vanA/B* | Mis-sing | *optrA* | *cfr(D)* | *optrA* + *cfr(D)* | G2576T | Mis-sing |
| ST117 | ST143 | 7799<sup>a</sup> | 2022-2024 | 198 | 223 | 21 | 156 | 43 | 3 | 5 | 4 | 62 | 2 | 148 |
|  | ST143 | 6173 | 2023 | 1 |  |  |  |  |  |  |  |  |  |  |
|  | ST18 | 929/2505 | 2023-2024 | 11 |  |  |  |  |  |  |  |  |  |  |
|  | ST128 | 118<sup>a</sup> | 2021-2024 | 5 |  |  |  |  |  |  |  |  |  |  |
|  | ST129 | 8952<sup>a</sup> | 2023-2024 | 6 |  |  |  |  |  |  |  |  |  |  |
|  | NA | NA | 2023-2024 | 2 |  |  |  |  |  |  |  |  |  |  |
| ST80 | ST152 | 8599<sup>a</sup> | 2023-2024 | 15 | 20 | 18 | 1 | 1 | 0 | 0 | 0 | 19 | 0 | 1 |
|  | ST148 | 8954 | 2023 | 2 |  |  |  |  |  |  |  |  |  |  |
|  | ST877 | 8551<sup>a</sup> | 2023-2024 | 2 |  |  |  |  |  |  |  |  |  |  |
|  | ST150 | 6660 | 2023 | 1 |  |  |  |  |  |  |  |  |  |  |
| ST1886 | ST1330 | 2290 | 2021 | 5 | 5 | 5 | 0 | 0 | 0 | 0 | 1 | 4 | 0 | 0 |
| Other<sup>b</sup> |  |  | 2023-2024 | 5 | 5 | 3 | 0 | 0 | 2 | 1 | 0 | 1 | 0 | 3 |
| NA |  |  | 2021 | 1 | 1 | 1 | 0 | 0 | 0 | 0 | 0 | 0 | 0 | 3 |
| Total |  |  |  |  | 254 | 48 | 157 | 44 | 5 | 6 | 5 | 86 | 2 | 155 |

ST: sequence type; cgMLST: core genome multilocus sequence typing; CT: complex type; NA: ST undetermined.
<sup>a</sup> Includes isolates for which CT could not be determined. The most common CT associated with each ST (Bezdiček) is shown.
<sup>b</sup> Includes one isolate each of ST17, ST217, ST327, ST494 and ST1478.

We characterized isolates harbouring the *vanA* operon (n=48, 18.9%), *vanB* operon (n=157, 61.8%) and *vanA*+*vanB* (n=44, 17.3%), with five (2%) isolates missing *van* genes (Table 2). Most *vanB* (n=153/157) or *vanA*+*vanB* (n=43/44) isolates belonged to the dominant ST117-CT7799 strain. In contrast, all *vanA* isolates belonged to non-ST117-CT7799 (Table S2). Acquired linezolid resistance determinants were identified in 99 isolates (38.9%). In most of these cases (n=86/99, 86.9%), the resistance genes *optrA* and *cfr(D)* were found together. Only two (<1%) unrelated isolates (ST117-CT929 and ST117-CT118) only had chromosomal G2576T mutations in 23S rRNA genes. Most isolates with *optrA* and/or *cfr(D)* (n=97) also carried *vanA* (n=45/97) or *vanA+vanB* (n=44/97) (Table 2, Table S2).

### Phylogenomic analysis

Phylogenetic reconstruction based on the cgSNPs of the soft core (95%) split ST80 and ST117 (Homan scheme) lineages, in contrast with strict core genome phylogeny (Figure S2). Moreover, VREfm were grouped in 7 main clades (including n>2 isolates), largely consistent with CTs and STs (Bezdíček scheme) (Figure 2).

**Figure 2.**
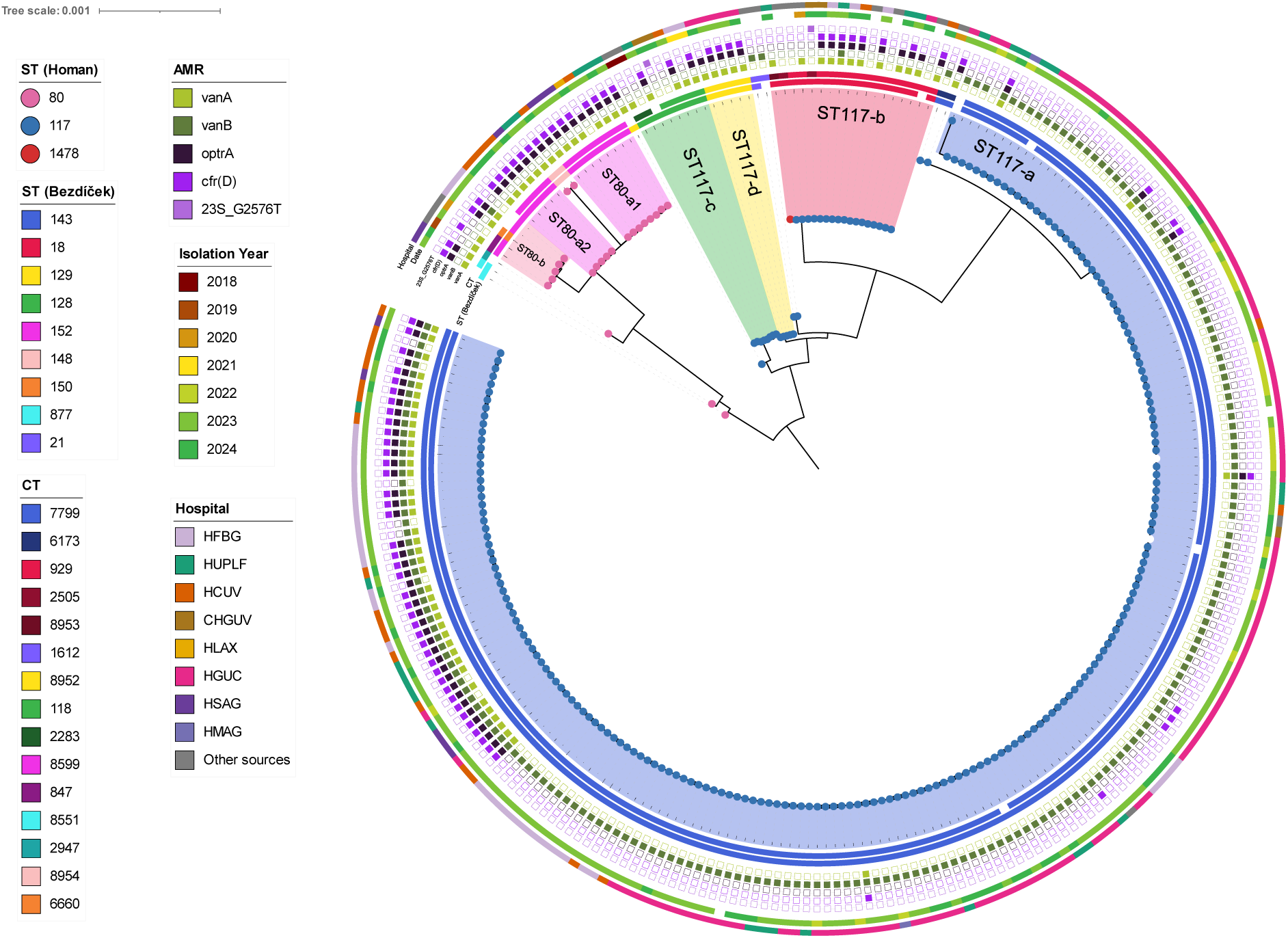
Maximum likelihood phylogeny of ST80 and ST117 Efm genomes from the Comunitat Valenciana, Spain. The final alignment had 2,363,951 relaxed core (95%) positions with 11,069 variable sites. The tree was rooted at midpoint and visualized using iToL v7. Internal nodes with bootstrap <95% are indicated with a dot. The closest sequences to each lineage from external sources are included within the tree.

ST117-CT7799 short-read sequences were grouped in a single phylogenetic clade (ST117-a, n=194) with a median pairwise distance of 9 SNPs (average 9.99, range 0-49). Furthermore, 192 isolates across seven hospitals had <10 SNP differences to one sequence within the same clade, suggesting transmission (Figure 3, Table S4). Despite high within-clade genomic similarity, CT7799 phylogeny was structured by ARG content and hospital, with most *vanA^+^/vanB*^+^, *optrA^+^*, *cfr(D)^+^* isolates found in a single subclade (Figure 4). Clades ST117-b (n=12) and ST117-c (n=5) mostly included closely related isolates (<10 SNPs), while ST117-d (n=5), which had the largest range of isolation dates (February 2021 - July 2024), also showed the largest intra-clade pairwise SNP distances among ST117 lineages (Figure 2, Figure 3, Table S4). In contrast, ST80 clades were more genetically diverse than ST117 clades (Figure 3). The only clades associated with a single hospital were ST117-c (HGUC) and a subclade of ST80-b (HFBG) (Figure 2).

**Figure 3.**
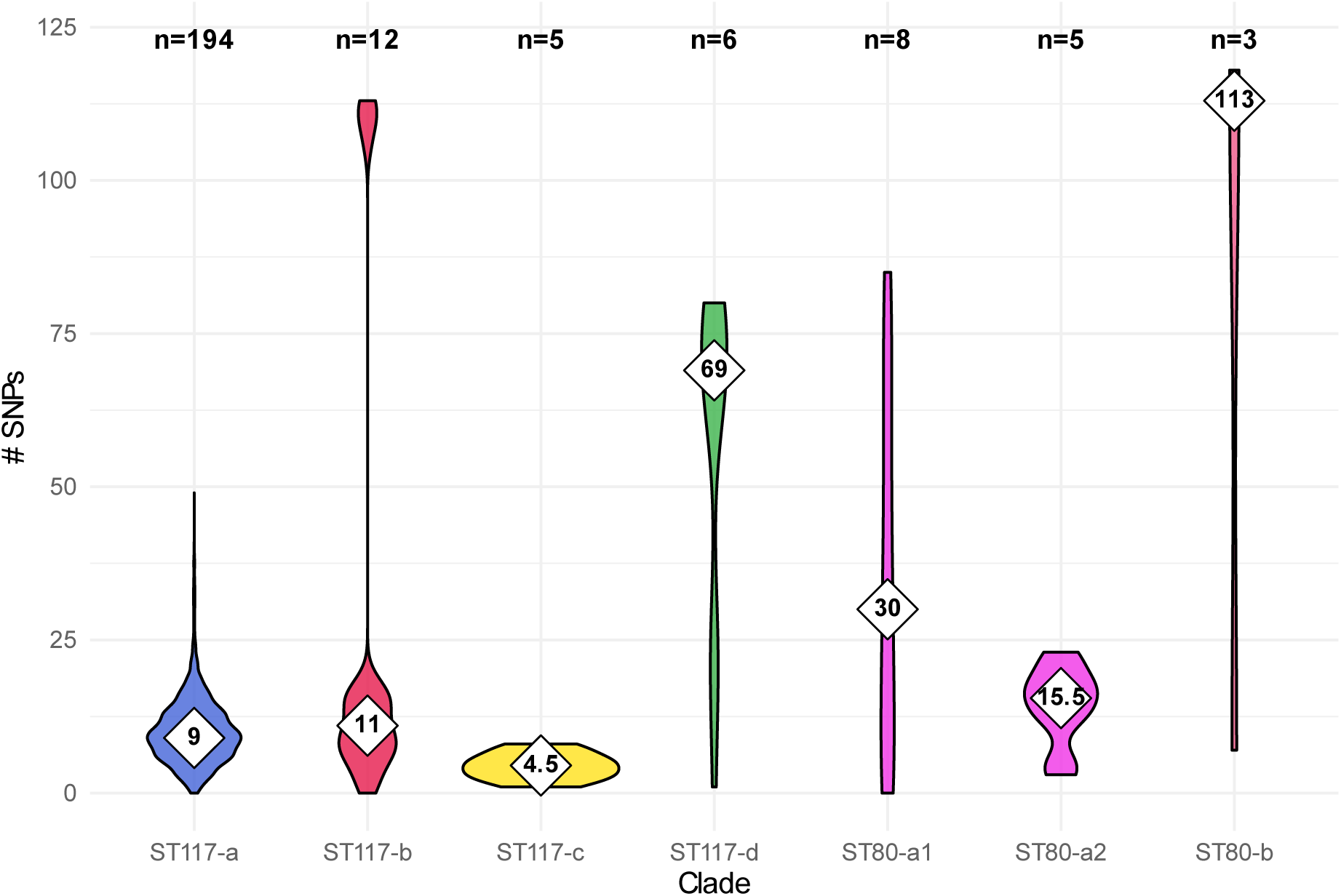
Distribution of intra-clade SNP distances. Only clades with n>2 genomes are shown. Median values are shown inside rhombus. The number of isolates compared are shown at the top of each violin plot. Only CV genomes were considered for the distance calculation.

**Figure 4.**
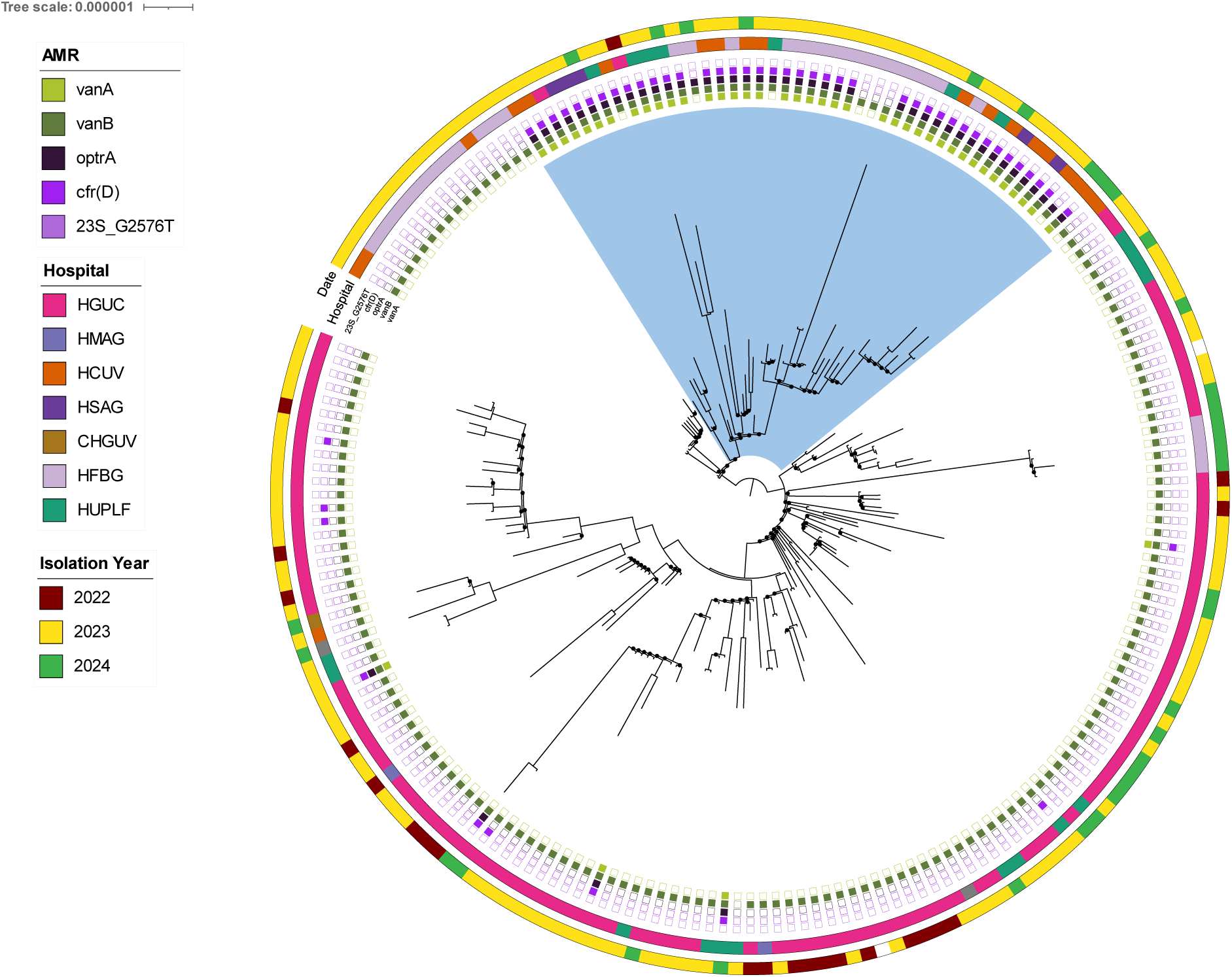
Maximum likelihood phylogeny of ST117-CT7799 clade. *vanA/B* clade is highlighted in blue. Alignment had 2,674,320 relaxed core (95%) positions with 364 variable sites. Tree was rooted at midpoint and visualized using iToL v7. Internal nodes with bootstrap <70 are indicated with a dot.

Some of the publicly available complete genome sequences from different sources were closely related to the CV lineages. Notably, two ST117-CT7799 isolates (2023-2024) from other regions in Spain clustered with the CV sequences, differing by <10 SNPS from five and 144 CV genomes, respectively (Figure 2, Table S3, Table S4). Furthermore, two ST80-CT847 strains isolated in Barcelona, Spain (2020 and 2023, respectively) formed a clade with ST80 CV isolates from different CTs. Sequences from Germany (n=3) and Denmark (n=2) grouped with ST117-CT929/CT2505. Interestingly, two strains isolated in Ireland in 2018 were included within the ST117-CT118 clade (Figure 2). Comparison with ST143 (Bezdíček scheme) complete genomes in ATB revealed that CT7799 CV samples constituted a separate clade. The median distance to the sister clade (CT6173), which included EPI00457 and closely related strains (Switzerland, 2020), was 270 SNPs (Figure S3).

### Antimicrobial resistance, virulence and bacteriocin profiles

Twenty-one different acquired ARGs were identified across the complete dataset, with most strains carrying between 3 and 10 acquired ARGs (Table S2, Table S5), excluding low quality sequences (Table S6). Apart from the *vanB* operon, the most frequent genes (n=186-239, 73-94%) conferred resistance to aminoglycosides-streptothricin (*aac(6’)-ie-aph(2’’)-Ia*, *ant(6)-Ia*, *aph(3’)-IIIa*, *sat4*), macrolides-lincosamides-streptogramin B (*ermB*) and trimethoprim (*dfrG*) (Table S5), and were broadly distributed across hospitals and lineages (Figure S4). Interestingly, *vanA*, *optrA*, *cfr(D)* and *ermA* were consistently found together across all *vanA^+^*and *vanA*+*vanB* clones. Less prevalent ARGs (*tetM, tetL*) were observed without a clear association with phylogeny, hospital or particular *rep* genes (Figure S4). Most isolates (n=251, 98.8%) carried between 1-4 daptomycin-associated mutations. Type and frequency of the mutations varied across ST-CTs, with most ST117 isolates harbouring *liaR_S19F, rpoC_T641K*, and *dltC_S63C*, or at least two of these mutations. In contrast, *cls_A20D* was found only in two ST117-CT7799 isolates. Furthermore, most ST80 isolates harboured *liaR_E75K*, *liaR_W73C* and *liaS_T120A*, while rare STs carried a single mutation (Table S5).

Thirty-six different PVMs were identified, with VREfm strains harbouring 18-34 PVMs (median 31), mostly corresponding *E. faecium* hospital gene variants (Table S2, Table S8). Additionally, seven genes were associated with specific clades (Figure S4). *capD* and *tirE2* were found exclusively in ST117, while PGC-1 (*fms20, fms21*) was predominant in non-CT7799 isolates. Interestingly, ST117-CT929 and related lineages CT2205 and CT8953 (ST117-b clade) exclusively harboured *tirE1* and a complete *ecbA* hospital gene variant. *fms15* and *scm* were variable, with the latter being more prevalent in CT7799 strains from one hospital (HFBG).

Nine different bacteriocin genes were identified, with 1-7 genes detected per strain (Table S2, Table S9). Most (n=251, 99%) carried *entA*, followed by *bacAS9* (87%), which was truncated in all but one case. All *bac43*-carrying isolates also harboured *bacAS9* (Table S7). The combination of these three genes was observed across all ST117 and ST80 lineages and was the most prevalent bacteriocin profile (n=186, 73%).

A total of 38 different replicases from six families were identified in the complete dataset, with strains carrying between 1-13. The most frequent plasmid families as represented by their *rep* genes were Rep_trans (98% strains), Rep3 (97%), RepA_N (97%) and Inc18 (83%) (Table S7).

### Analysis of complete plasmids

Using hybrid and long-read assembly, we generated complete assemblies of 38 *E. faecium* genomes (30 *vanB*, 6 *vanA*, 2 *vanA*+*vanB*; total assembly size: 2.89-3.34 Mb; Table S20) representing four hospitals (2022-2024). These isolates harboured 193 plasmids (3-7 per strain), ranging from 1.9 to 293 kb. The *vanB* operon was located in the same chromosomal region in all 32 *vanB*-only and *vanA+vanB* strains, while the *vanA* operon was always carried on linear plasmids (117-121 kb) in the eight *vanA-*only and *vanA+vanB* strains. All linear plasmids consistently carried three Rep3 replicases (rep_US78__pZY2, rep_US80__pSRR24, rep_41__PR02395-7p2) and co-harboured *vanA, optrA, cfr(D)* and *ermA*.

Furthermore, all isolates carried Rep3 or Rep_trans small plasmids (2-11 kb, 113/193), which showed the largest diversity in *rep* genes. Rep3-like plasmids with rep_11a__pB82 (6 kb) and rep_44__M1215241p4 (4 kb) exclusively carried *bac43* and *bac51* bacteriocins, respectively. Medium-sized plasmids (28-39 kb, 34/193) carried an identical replicase (Inc18::rep_2__pRE25) and ARGs [*aph(3’)-IIIa, ermB, SAT-4, aad(6)*]. Megaplasmids (162-293 kb, 37/193) had different RepA_N replicases in CT7799 (rep_US15__pNB2354p1) and non-CT7799 (rep_US15__DO3), although consistently harboured common ARGs [*aac(6’)-Ie-aph(2’’)-Ia, aph(3’)-IIIa, ermB, SAT-4, aad(6)*] and PVMs (*hyl,* PGC-1 cluster).

The eight *vanA* linear plasmids grouped in a single community using the pling tool (Figure 5b), showing high sequence (containment 0.00-0.03) and structural similarity (DCJ-indel 0-4) (Figure 5a). Horizontal transfer of these nearly identical plasmids is suggested from the phylogeny and hospital distribution, as sequences were recovered from eight strains belonging to six different clones within ST17, ST80 and ST117 (ST117-CT7799, ST117-CT118, ST117-CT929, ST80-CT8599, ST80-CT8551 and ST17-CT8951) and across four hospitals (HCUV, HFBG, HGUC, HSAG) during 2022-2024 (Figure 5b, Table S20).

**Figure 5.**
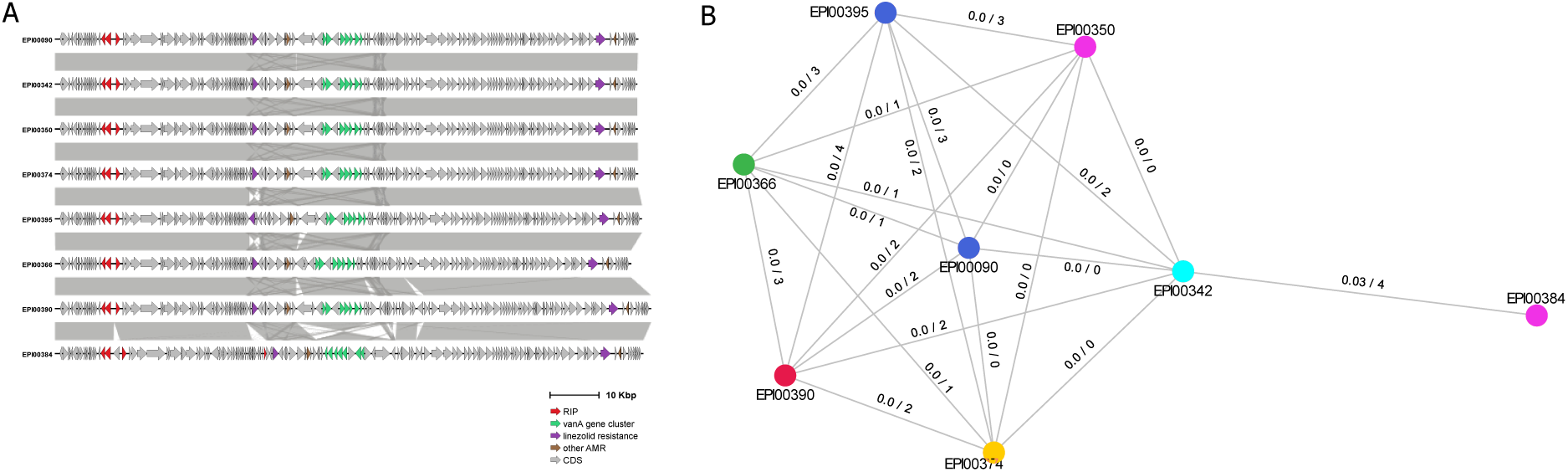
Comparison of *vanA* linear plasmids. (A) Comparison of plasmid sequences. CDS are represented as arrows and links are shown when sequence similarity is >99%. (B) Clustering of *vanA* plasmid sequences (nodes) with pling, showing containment/DCJ-indel distances at edges. Nodes are coloured by ST and are connected if containment distance is ≤0.3.

## DISCUSSION

This retrospective genomic study provides the first region-wide investigation of inter-hospital spread of VREfm in the Comunitat Valenciana, Spain. We describe (i) widespread circulation of multiple lineages across hospitals, (ii) marked clonal expansion of ST117-CT7799 during 2022–2024, and (iii) horizontal spread of Rep3-like linear plasmids co-carrying clinically critical resistance determinants, including *vanA, optrA* and *cfr(D)*. Together, these findings indicate that both clonal transmission and plasmid-mediated convergence are shaping the regional VREfm epidemiology and may facilitate the emergence of difficult-to-treat multidrug-resistant infections.

Multiple VREfm clones were detected across CV hospitals, dominated by the globally distributed ST117 and ST80 backgrounds. Although several CTs are seldom reported in clinical or outbreak settings (e.g., ST117-CT118 [36]), comparison with publicly available genomes revealed closely related isolates from other European countries (2018-2023), suggesting broader dissemination and/or repeated introductions. Notably, the placement of vancomycin-susceptible ST117-CT929 and ST117-CT2505 isolates within the CV ST117 diversity is consistent with recent reports describing these lineages in Germany [37]. In defining circulating lineages, Bezdíček’s *et al*. MLST scheme [18], still rarely applied in epidemiological studies [38], showed good concordance with phylogeny and cgMLST in our dataset, supporting its utility for surveillance.

The dominant ST117-CT7799 VREfm clone has only been sporadically described in clinical contexts in Spain [5]. Our phylogenomic analysis including publicly available genomes is consistent with a recent, substantial expansion in CV and suggests interregional spread within Spain during 2022-2024. The predominance of *vanB* within CT7799 differs from the broader Spanish pattern, where *vanA* has been more frequent [39] and large hospital outbreaks attributed specifically to ST117 *vanB E. faecium* have been reported only rarely [4]

However, ongoing shifts in the relative frequency of *vanA, vanB,* and *vanA+vanB* genotypes have been documented across Europe in recent years [7,39]. Beyond glycopeptide resistance, nearly 20% of CT7799 and most ST80 isolates harboured *optrA* and *cfr(D)*, in line with the global increase of LREfm/LVREfm infections in recent years [40,41] and contrasting with earlier studies in which 23S rRNA mutations predominated among linezolid-resistance mechanisms [12,42]. Importantly, in Spain these genes were largely confined to *E. faecalis* in hospitals before 2020 [43,44], highlighting the public health relevance of their emergence in *E. faecium* lineages with clear evidence of hospital spread.

Regarding daptomycin resistance, ST117-CT7799 frequently harboured mutations in the *liaFSR* operon, an established pathway to reduced daptomycin susceptibility in *E. faecium*, often alongside additional secondary/compensatory mutations in *rpoC* and *dltC* that may contribute to resistance and/or fitness. In contrast, combinations of *liaFSR* and *cls* mutations, which have been associated with higher-level resistance, were uncommon [45–47]. However, because minimum inhibitory concentration (MIC) data were not available, genotype–phenotype interpretation for daptomycin should be made cautiously [48].

Hybrid assemblies enabled detailed characterization of a Rep3-like linear plasmid co-harbouring *vanA, optrA* and *cfr(D)* across multiple clones and hospitals, strongly suggesting ongoing plasmid transmission. Reports of linear plasmids in *E. faecium* have increased in parallel with the use of long-read sequencing, which is essential to resolve plasmid structure and topology [49,50]. The co-localization of these three ARGs on enterococcal linear plasmids has recently been reported in India, Italy and Canada [35,51,52], and related *vanA* plasmid backbones (e.g., rep_US78__pZY2) appear to circulate internationally [34]. In our dataset, acquisition of this linear plasmid by a CT7799 *vanB* subclade provides a plausible explanation for the notable proportion of *vanA*+*vanB* isolates (>17%), a genotype generally considered uncommon in VREfm collections [7].

Finally, CT7799 consistently carried a small *bac43*-positive plasmid (rep_11a__pB82-like; 6,173 bp) previously described internationally [23,53]. The increasing prevalence of plasmid-borne bacteriocins such as *bac43* has been proposed to promote hospital adaptation and may contribute to lineage replacement and outbreak potential [23,53]. Conversely, plasmid-borne *bac51* was strongly associated with *vanA* isolates, yet its detection in CT7799/*vanB* suggests inter-lineage exchange of accessory traits, reinforcing the view that plasmids are key drivers of ecological success in hospital-adapted *E. faecium*.

This study has limitations. The retrospective design and lack of patient-level epidemiological data preclude formal reconstruction of transmission chains and directionality of spread. In addition, the availability of public genomes is uneven across regions and time, which may bias inferences on geographic dissemination. A more intense effort to obtain and make publicly available WGS of VREfm isolates would greatly facilitate a better understanding of the transmission and spread routes. Finally, the absence of phenotypic susceptibility results limits the clinical interpretation of some resistance-associated mutations. Still, our findings have immediate public health relevance, as the convergence of vancomycin and linezolid resistance in hospital-adapted *E. faecium* heightens the risk of hard-to-treat outbreaks and underscores the need for strengthened screening and genomic surveillance, including long-read sequencing to accurately track linear plasmids. Expanded genomic surveillance, ideally coupled to epidemiological and antimicrobial susceptibility data and including long-read sequencing for plasmid resolution, would improve understanding of transmission routes and inform control measures.

## CONCLUSIONS

This study reports the inter-hospital dissemination of VREfm lineages in the Comunitat Valenciana during 2022-2024, with genomic links to other Spanish regions and Europe. Our data evidences a marked regional expansion of ST117-CT7799 and reveal horizontal transfer of a Rep3-like linear plasmid co-carrying *vanA*, *optrA* and *cfr(D)*, increasing the risk of multidrug resistance and therapeutic failure. These findings highlight the need for strengthened infection prevention and control and highlight the added value of genomic surveillance, particularly long-read–enabled plasmid reconstruction, to detect and interrupt transmission of high-risk VREfm lineages and mobile resistance elements.

## Statements

## Data availability

All sequence reads have been submitted to ENA under the project number PRJEB110923.

## Authors’ contributions

CVM: methodology, formal analysis, visualisation, manuscript drafting. ACAS: methodology, data interpretation, manuscript review and editing. LRR: genome sequencing, investigation, conceptualization, manuscript review and editing. RA: genome sequencing. AS: software, data curation. SA: genome sequencing, methodology. SS, IT, EGB, IV, NT, RF: sample collection and screening of the isolates. CN, LP: conceptualization, data interpretation. ARF: conceptualisation, methodology, data interpretation, manuscript review and editing. FGC: conceptualisation, methodology, data interpretation, manuscript review and editing, supervision.

## Ethical statement

Ethical approval was not required for this study as only data from routine surveillance was included.

## Use of artificial intelligence tools

None declared.

