## Supplemental Figures S1-S5 for "Multihospital expansion of vancomycin-resistant *Enterococcus faecium* ST117-CT7799 and transmission of linear plasmids co-carrying *vanA* and linezolid resistance genes, Comunitat Valenciana, Spain (2022–2024)"

### SUPPLEMENTARY MATERIAL

**Figure S1.** Distribution of the hospitals included in the study.

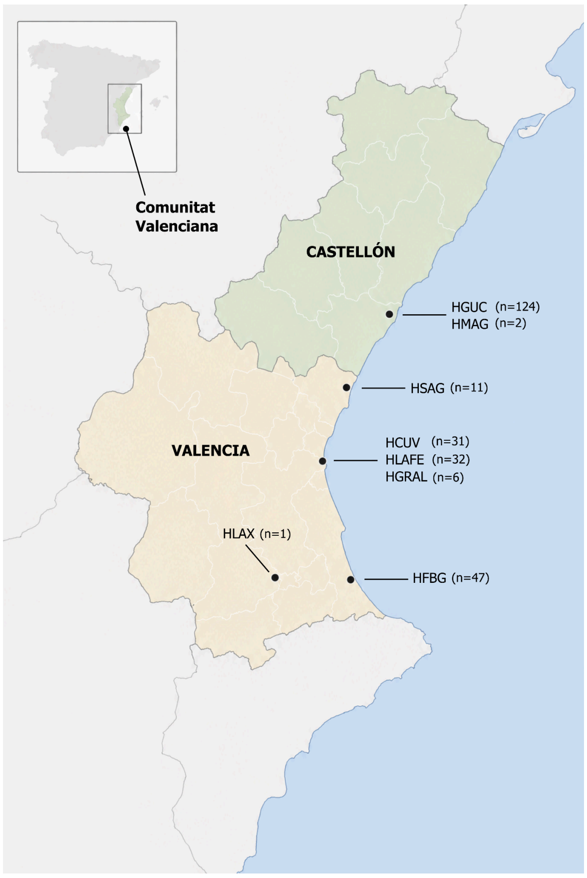


HCUV: Hospital Clínico Universitario de Valencia; HFBG: Hospital Francesc de Borja de Gandía; HGRAL: Consorcio Hospital General Universitario de Valencia; HGUC: Hospital General Universitario de Castellón; HLAFE: Hospital Universitario y Politécnico La Fe; HLAX: Hospital Lluís Alcanyís de Xàtiva; HMAG: Hospital de la Magdalena; HSAG: Hospital de Sagunto.

**Figure S2.** Maximum likelihood phylogenies of CV *E. faecium* genomes including rare STs.

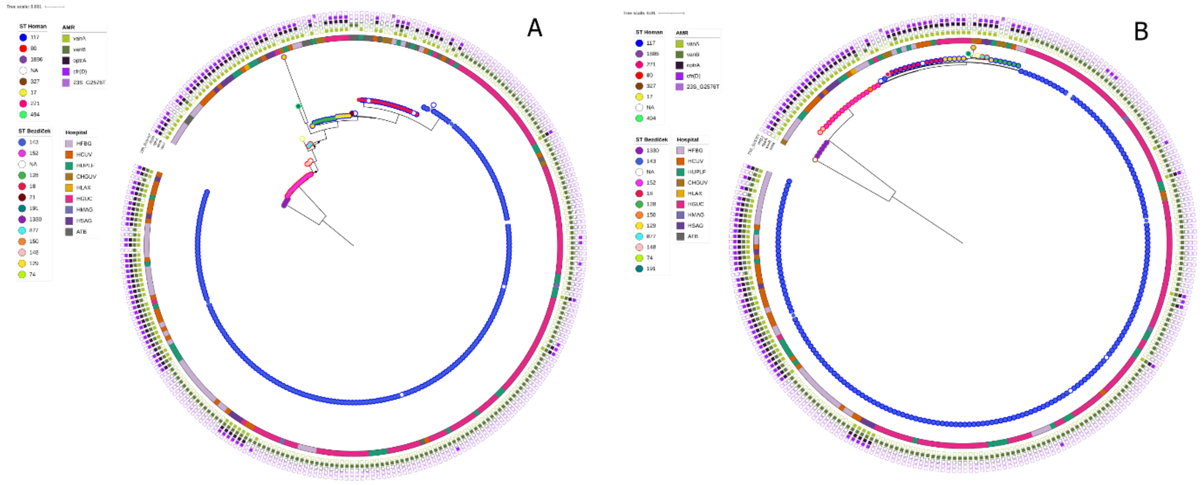


Alignments had 2,303,793 relaxed core (95%) positions with 113,330 variable sites. Trees are rooted at midpoint. Internal nodes with bootstrap <90 are indicated with a dot. The color of the border of each isolate's point represents Homan scheme's ST, while the internal color represents Bezdicek scheme's ST. (A) ML tree including rare STs except 1 ST327 isolate for better visualization. (B) ML tree including all isolates from rare STs. Trees were visualized using iToL v7.

**Figure S3.** Maximum likelihood tree from strict core of ST80 and ST117 *E. faecium* genomes.

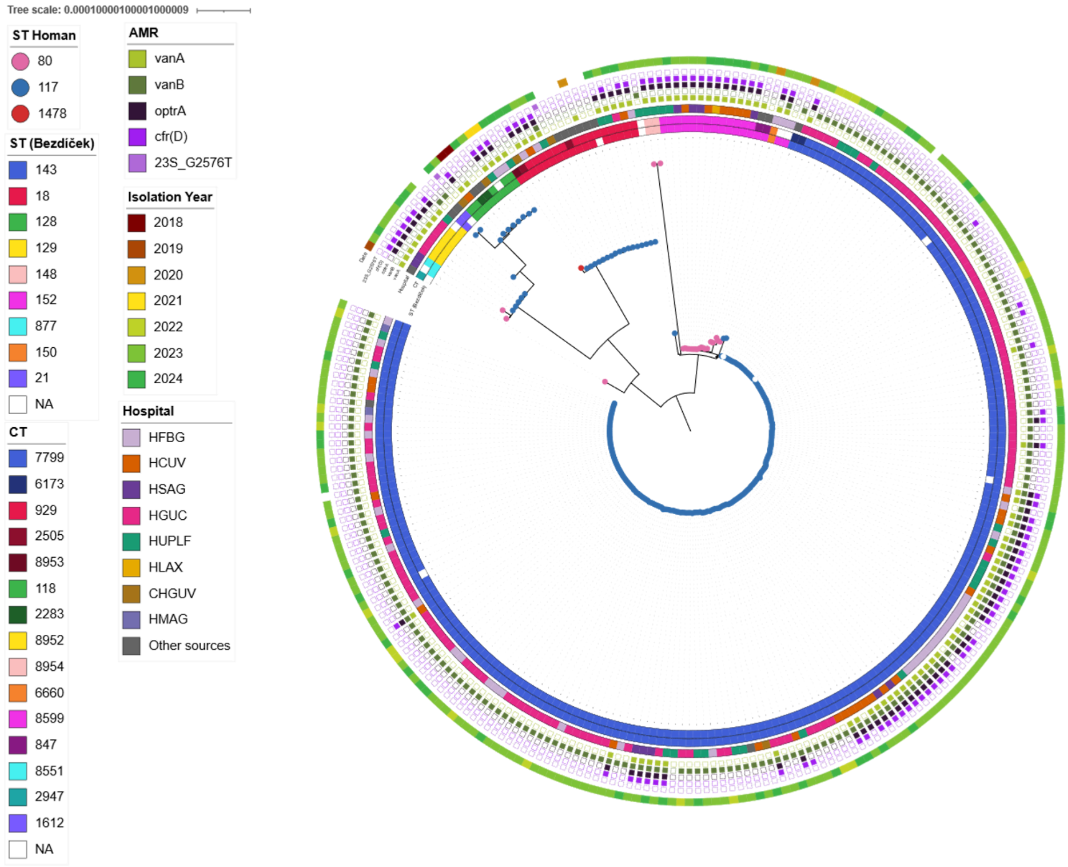


Alignment had 2,879,387 positions with 3,181 variable sites. Isolation location is indicated only for the sequences from external sources. The closest sequences to each lineage from external sources are included. Tree was was rooted at midpoint and visualized using iToL v7.

**Figure S4.** Maximum likelihood tree of ST143 *E. faecium* (Bezdíček scheme).

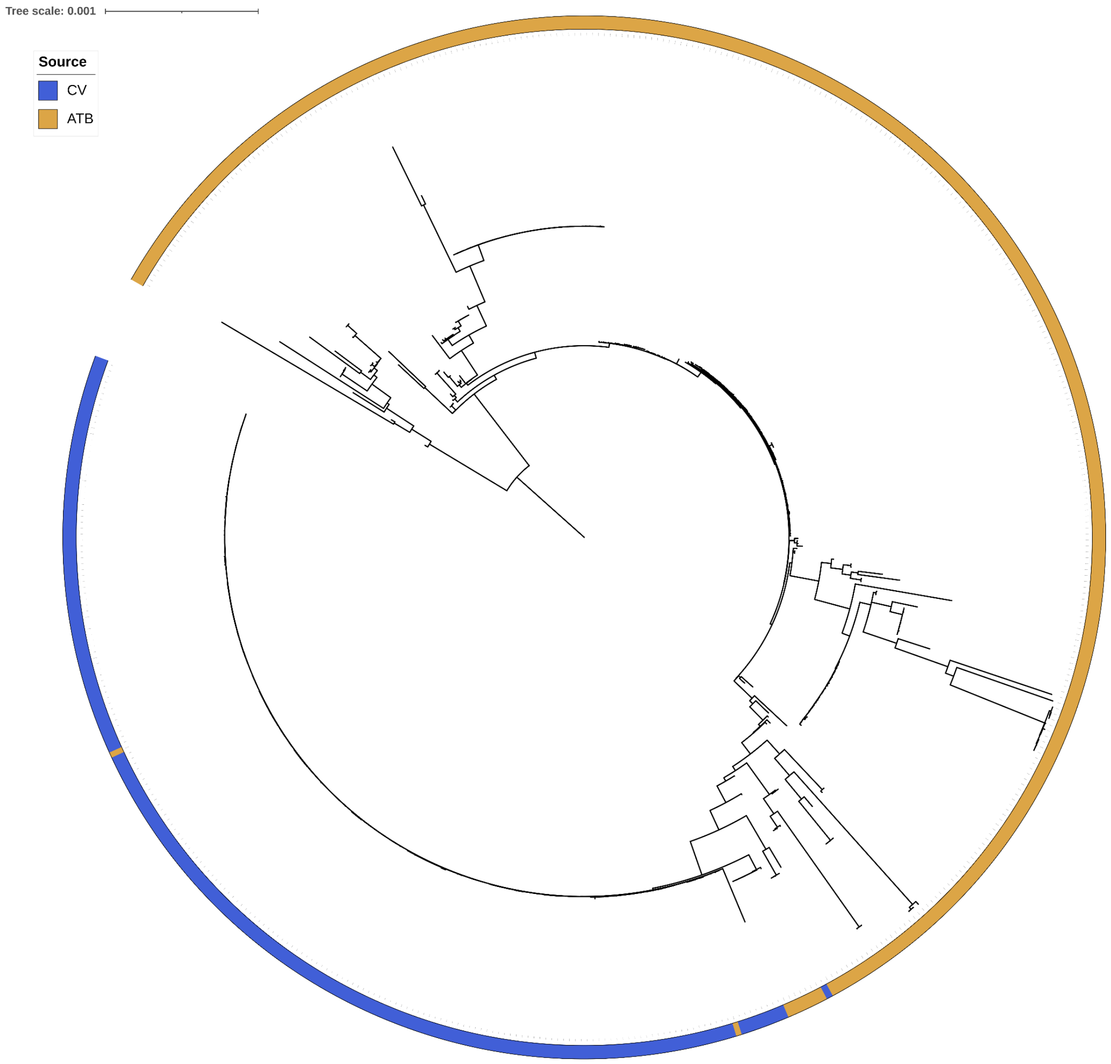


Tree includes ST143 (Bezdíček scheme) *E. faecium* genomes from this study (CV), AllTheBacteria database (ATB) and Pitart et al. [5]. Alignment had 2,182,943 relaxed core (95%) positions with 17,099 variable sites. Tree was was rooted at midpoint and visualized using iToL v7.

**Figure S5.** Distribution of *rep* genes, bacteriocins, PVM and ARGs in the phylogeny of the complete dataset.

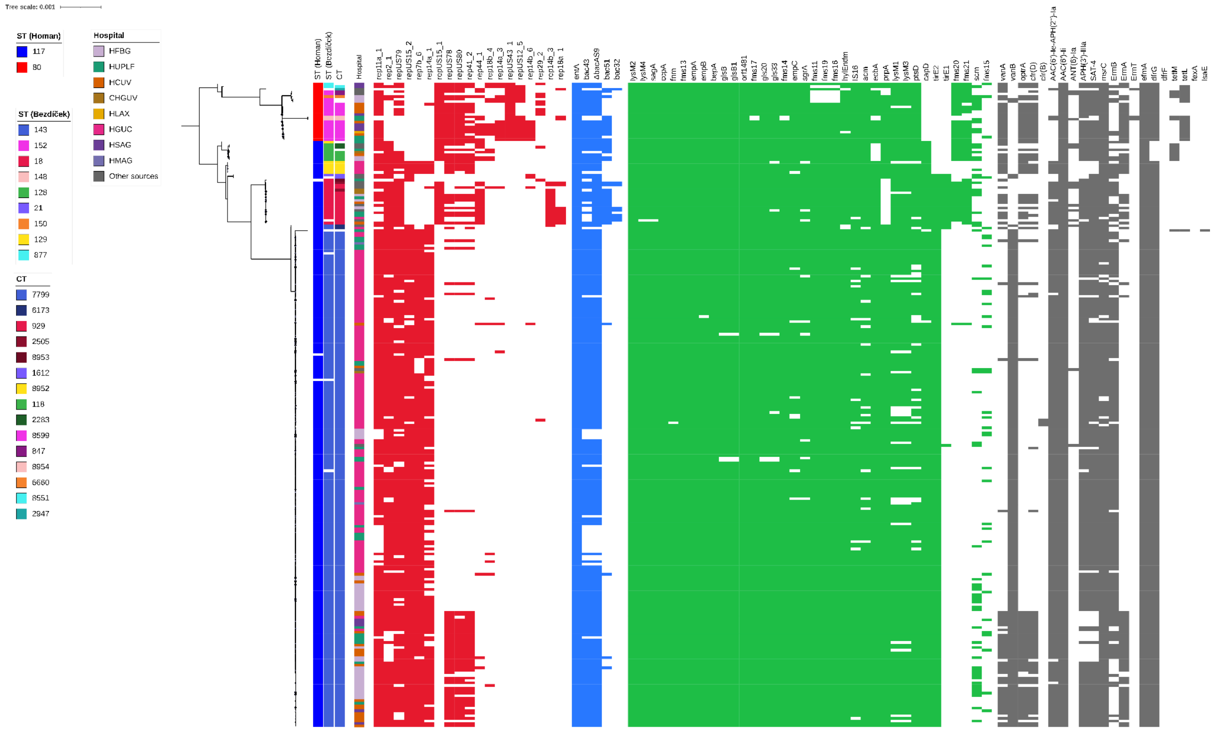


Presence of rep genes, bacteriocins, PVM and ARGs is represented by red, blue, green and grey squares, respectively. ML phylogeny of ST80 and ST117 genomes was built from relaxed core (95%) alignment (2,363,951 positions, 11,069 variable sites). Tree was was rooted at midpoint and visualized using iToL v7. Internal nodes with bootstrap <95 are indicated with a dot.
